# Hybrid Transformer and Neural Network Configuration for Protein Classification Using Amino Acids

**DOI:** 10.1101/2024.10.31.621421

**Authors:** Nathan Labiosa, Aryan Kohli, Christian Chung, Christopher Korban

## Abstract

This study introduces a hybrid machine learning model for classifying proteins, developed to address the complexities of protein sequence and structural analysis. Utilizing an architecture that combines a lightweight transformer with a concurrent neural network, the hybrid model leverages both sequential and intrinsic physical properties of proteins. Trained on a comprehensive dataset from the Research Collaboratory for Structural Bioinformatics Protein Data Bank, the model demonstrates a classification accuracy of 95%, outperforming existing methods by at least 15%.

The high accuracy achieved demonstrates the potential of this approach to innovate protein classification, facilitating advancements in drug discovery and the development of personalized medicine. By enabling precise protein function prediction, the hybrid model allows for specialized strategies in therapeutic targeting and the exploration of protein dynamics in biological systems.

Future work will focus on enhancing the model’s generalizability across diverse datasets and exploring the integration of more machine learning techniques to refine predictive capabilities further. The implications of this research offer potential breakthroughs in biomedical research and the broader field of protein engineering.

## I. Introduction

Proteins perform many roles in biological systems and their amino acid sequences dictate their structural conformations and function. For instance, proteins can function as enzymes, provide structural support to cells, facilitate material transport, serve as therapeutic agents like monoclonal antibodies, participate in DNA replication and transcription, and act as precursors for other protein formations. Further exploring categorization, proteins can be classified into several types, including signaling proteins, immune system proteins, and transcription proteins, etc. Additionally, enzymes are categorized into six principal groups: oxidoreductases, transferases, hydrolases, lyases, isomerases, and ligases. This paper introduces a novel computational approach to classify a vast array of protein types, leveraging only the primary structure of the protein and ten intrinsic properties.

Traditional protein classification relies on a group of sophisticated laboratory techniques. Gel electrophoresis, for instance, separates peptides by molecular weight and uses staining methods such as Coomassie blue and silver stain for visualization. Complementing this, structural assessments frequently apply X-ray crystallography and electron microscopy to examine protein formations. Another technique, Static Light Scattering, determines molecular weight, solubility, and aggregation propensity through light scattering metrics. Further, methods like mass spectrometry and isoelectric help to enhance the accuracy of classifying distinct protein types. Despite their precision, these methods are often time-consuming and costly [1]. With the roadblock of extensive wet-lab data, there arises a substantial opportunity for efficiently applying machine learning to enhance protein classification. Particularly, if machine learning models can swiftly classify proteins, it could potentially speed up evaluating protein function up to a hundred times faster than traditional techniques [2].

The significance of rapid protein classification extends to several areas. Primarily, it drives the assessment of protein functions, which is crucial in the discovery and development of biologics. Efficient protein classification not only facilitates the examination of potential therapeutic impacts but also allows for the study of functional changes resulting from amino acid mutations. In the fields of proteomics and drug repurposing, accurate protein identification and expression level analysis are key steps towards discovering new therapeutic targets [3]. This combined approach holds promise for expediting the identification and utilization of novel protein targets in drug development research.

Despite the deep knowledge base in the protein field, the demand for continued research in proteomics and biologics, particularly for personalized medicine, remains strong [4]. With the protein sector projected to grow at a compound annual growth rate of 15.53% through 2030, [5], there is a need for innovative solutions to keep pace with this expansion. In response, machine learning has emerged as a powerful tool to propel advancements. For instance, generative models like AlphaFold 2 leverage deep learning for high-accuracy structural predictions, while ProtBert addresses evolutionary tasks and sequence masking. Further, efforts to engineer new amino acid sequences use advanced techniques such as generative adversarial networks and variational autoencoders. On the discriminatory side, models like random forests and logistic regression are used to predict important physicochemical properties of proteins, including pH, water solubility, and thermal stability. Additionally, classification models are increasingly being deployed to identify subcellular localizations and facilitate protein identifications. Despite the potential of these methods, the increasing complexity and scope of protein research emphasizes a critical need for superior accuracy in protein identification tasks [3].

Recent advancements in natural language processing have catalyzed the development of new architectures, notably the transformer model. This architecture utilizes a mechanism of query, key, and value pairs, enabling the model to selectively ‘pay attention’ to different sequences within the input [6], [7]. This attention mechanism allows the model to adjust the significance it assigns to each part of the input based on its context, significantly enhancing the model’s ability to interpret complex data sequences [8], [9], [10]. The versatility of transformers has led to their widespread application across various domains, including the creation of expansive language models for text analysis and hybrid models that integrate computer vision with linguistic processing [11], [12], [13]. The attention mechanism of transformers holds immense promise for biological applications, particularly in the study of amino acids. In the context of proteomics, each amino acid sequence correlates directly to a physical molecule, with the sequence order fundamentally changing the molecule’s properties. By harnessing the attention mechanism, it becomes possible to analyze and classify amino acids based on their sequential characteristics. This capability is key for identifying and categorizing new amino acid sequences as specific protein types. Successfully classifying proteins through their amino acid sequences can revolutionize researcher’s understanding and manipulation of biological functions. It could lead to breakthroughs in designing novel proteins with desired properties, enhancing drug discovery, and tailoring therapeutic interventions more precisely to individual genetic profiles [14]. This could significantly impact the fields of personalized medicine and biotechnology, where accurate protein function prediction enables more targeted treatment strategies and the development of new biologic drugs [15].

In this paper, the following key contributions are presented:

- The introduction of an unique method that utilizes context-dependent machine learning tools to leverage both the sequential and physical properties of amino acids for classification.
- The details of the construction of a novel model architecture, designed to effectively integrate and process these dual features, resulting in a system capable of highaccuracy protein classification.

## II. Related Works

The application of transformer models to protein analysis has been attempted; significant among these attempts is the development of ProtBERT [16], [17]. ProtBERT was trained on over 106 million protein sequences using a bidirectional encoding strategy. This training involves masking specific amino acids within a sequence and tasking the model with predicting these masked positions, thereby enabling it to learn the intricate relationships between amino acid patterns [17]. While ProtBERT demonstrates the capabilities of transformers in biological contexts, it is primarily optimized for tasks other than protein classification. Furthermore, its training, though less extensive than some of its counterparts, required four weeks of pre-training on significant computational resources, rendering it impractical for researchers with limited access to time and GPU capabilities to duplicate [17].

Another noteworthy model in the field is ProtTrans [18]. Like ProtBERT, ProtTrans was primarily developed for predicting protein sequencing and structural configurations. The ProtT5-XL variant of this model underwent training on a supercomputer, utilizing thousands of GPUs and TPUs. Stemming from its immense computational demand, ProtT5-XL’s architecture is too large for deployment on standard consumer GPUs and similar to ProtBERT, it was not tailored for direct protein classification tasks [19].

Efforts to classify proteins using other machine learning methods have been documented, particularly applied to the same dataset that was utilized in this study. Notable among these are models employing Long Short-Term Memory networks, convolutional neural networks, and Naive Bayes classifiers [20], [21], [22]. These models leverage a combination of deep learning and statistical techniques to approach protein classification. Of these, the Naive Bayes model has shown the most success, achieving approximately an 80% accuracy rate in classifying proteins into 30 distinct classes.

## III. Method

The primary dataset features, aside from those derived computationally, were sourced from the Research Collaboratory for Structural Bioinformatics (RCSB) Protein Data Bank (PDB). The RCSB PDB is an open-source database in the U.S., offering structural data points for proteins, DNA, and RNA. All entries in the RCSB PDB undergo a careful review process, including consistency checks, cross-validation with other data sources, and verification with relevant literature [23].

Given the dataset volume, several preprocessing steps were necessary to prepare the data for the hybrid model. Initially, an analysis was conducted to determine an appropriate sequence length cutoff. With some sequences extending beyond 16,000 amino acids and the mean and median lengths around 350, this number was chosen as the cutoff to manage dataset variability Subsequently, the thirty most prevalent proteins were selected for inclusion in the model. This selection criterion was based on each protein having over 1,000 examples within the dataset, ensuring a significant sample size and reflecting the proteins’ regularity in biological research 3. Additionally, any duplicate sequences were excluded from the dataset.

To counteract model bias, which simpler preliminary models indicated was skewed towards the five most prevalent classes, a strategy of sample over-selection was employed. The Random Over Sampler technique was utilized to balance the class distribution within the training dataset [24]. This method works by randomly duplicating examples in the underrepresented classes until all classes have a comparable number of samples. This approach not only helps in reducing the model’s bias towards more frequent classes but also enhances the generalizability of the model across less common protein types.

To fully leverage the data available from protein analyses, amino acid sequences were utilized to extract additional physical and numerical features. The dataset initially included intrinsic features such as structureMolecularWeight, crystallizationTemperature (in Kelvin), Matthews coefficient (densityMatthews), percent solubility (densityPercentSol), and pH value. To augment this data, Biopython toolkit was employed, a tool for computational biology and bioinformatics. Biopython uses various biological computations, including the manipulation of protein sequences to extract biochemical properties [25]. A diagram of this extraction is shown in Figure 4.

From these sequences, critical features were extracted that are important in protein classification due to their impact on protein structure and function:

- Isoelectric Point: The pH at which a protein carries no net electric charge, significant for understanding protein solubility and interaction.
- Aromaticity: A measure of the frequency of aromatic amino acids, important for protein stability and function.
- Instability Index: A predictor of the stability of a protein in a test tube, with higher values indicating less stability.
- Helix, Turn, Sheet: Structural components of proteins, with helices and sheets forming the core structures and turns facilitating turns between them. These elements are necessary for defining the protein’s 3D conformation.

To ensure uniformity and improve the model’s performance, all numerical features were normalized. Additionally, protein sequences were front-padded to align them for processing by the model.

Building upon the integration of both sequential and numerical features, a novel model architecture was developed, as depicted in Figure 1. Initially, the study aimed to maximize the utility of amino acids represented as character sequences. To achieve this, a lightweight transformer architecture was employed designed to capture bidirectional relationships within the data. This choice was determined on the understanding that specific amino acids often neighbor each other, and their sequential arrangement influences protein function and classification [26]. The transformer’s ability to focus on different parts of the sequence makes it suited for tasks where context significantly affects the output, which is the foundation of protein classification.

**Fig 1:**
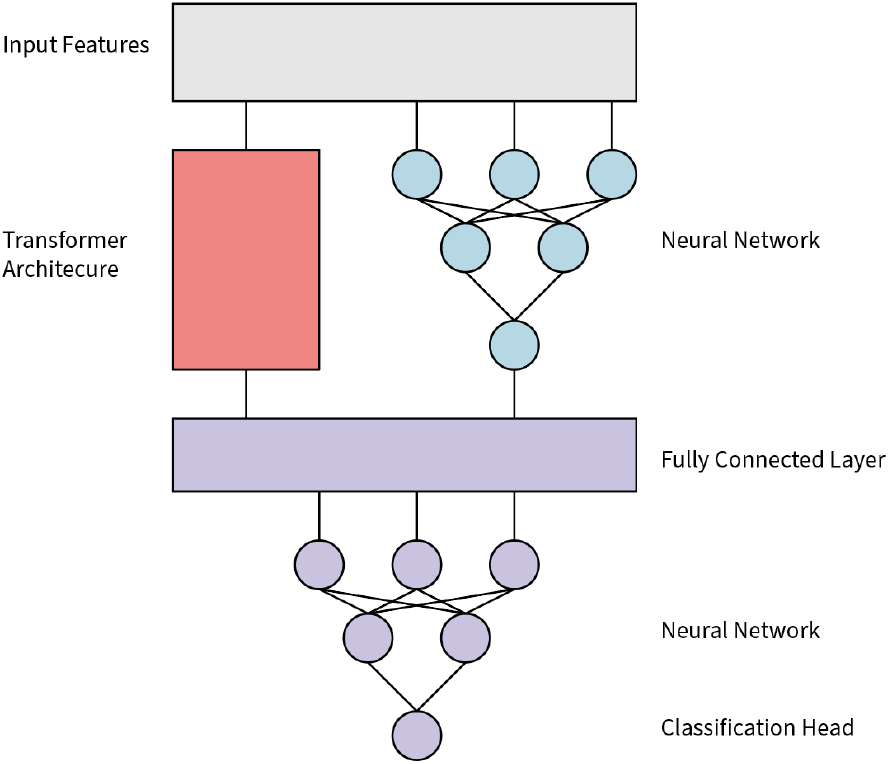
Protein Model Diagram

**Fig 2:**
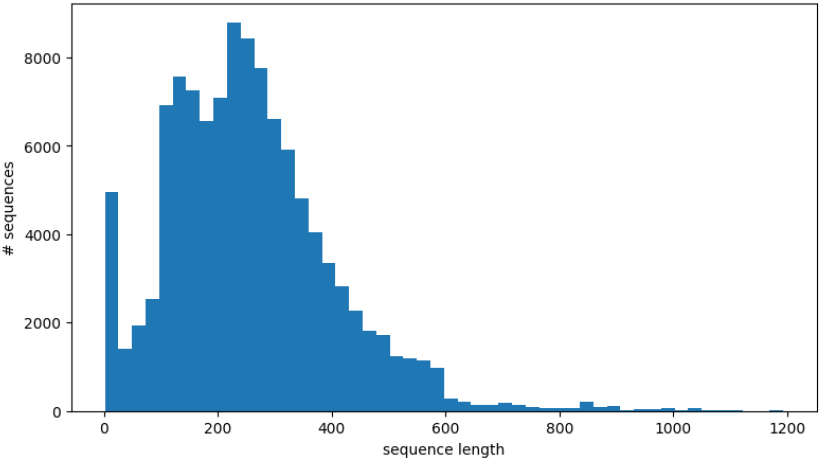
Sequence Length

**Fig 3:**
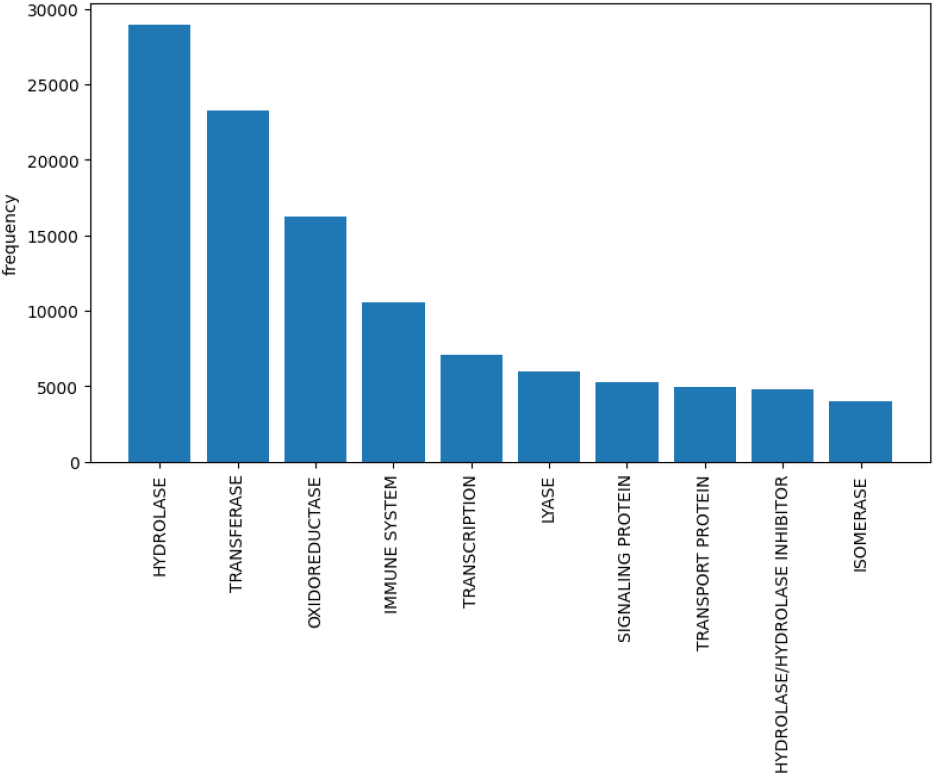
Class Distribution

**Fig 4:**
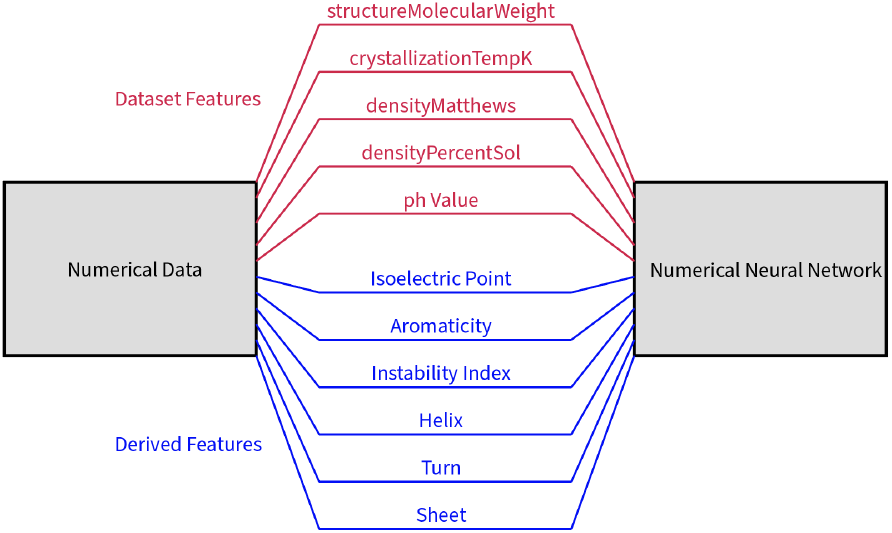
Numerical Data Diagram

Simultaneously, a neural network was engineered, tasked with processing the numerical features. This network was designed to interpret the physical properties of proteins, identifying key relationships within the latent space. The outputs from both the transformer and the concurrent neural network were then converged into a fully connected layer. This layer acts as a fusion point that combines learned sequential and physical insights, subsequently feeding into a neural network that results in a classification output.

This architecture represents a novel approach in protein classification, leveraging dual insights from amino acid sequences and their intrinsic physical properties. The combination of the sequential understanding provided by the transformer and the contextual insights from the numerical data processing allows for a more comprehensive classification. This method not only enhances accuracy but also offers a more holistic view of protein functionalities.

The model underwent training on Kaggle, utilizing their free, accessible GPU resources. The training process spanned approximately 10 hours, during which the model completed 13 epochs. This duration was chosen as it marked the point of diminishing returns, where further training ceased to yield significant improvements in model performance. The decision to halt training at this stage ensured the optimal use of computational resources while preventing overfitting.

## IV. Experimental Study

The outcomes of the study are detailed in Table I, where this novel model was benchmarked against previously mentioned protein classification methods on the PDB dataset. This study’s model achieves a classification accuracy of 95%, which is an improvement of at least 15% over the next best-reported technique. This increase in accuracy highlights the effectiveness of this novel approach.

**Table I:** * Indicates classification on only 10 classes. Otherwise the models performed classification on 30 classes.

| Method | Classification Accuracy (%) |
| --- | --- |
| Deep Neural Net * | 60 |
| LSTM * | 89 |
| AdaBoost | 18 |
| Transformer alone | 45 |
| Naive Bayes | 76 |
| Hybrid Approach | **95** |

Further demonstrating the robustness of the model, the evaluation metrics were expanded to include top 3 and top 5 accuracies. The results of these metrics are presented in Table II.This broader accuracy measurement further illustrates the model’s capability.

**Table II:** Model accuracy for top N predictions.

| Method | Protein Model Accuracy (%) |
| --- | --- |
| Top 1 | 95 |
| Top 3 | 99 |
| Top 5 | 100 |
| Top 10 | 100 |

To provide a practical idea of the model’s application, a detailed operation in a typical use case was documented. The process begins with extracting the amino acid sequences and initial numerical data from the dataset. Additional features are then derived using Biopython. This data undergoes normalization before being processed by the model, which then accurately outputs the protein’s class.

## V. Conclusion

This study introduced a novel approach to protein classification, leveraging a unique architecture that combines the strengths of transformer models and neural networks. By combining sequential and numerical features of proteins, this novel model achieved a classification accuracy of 95%. This result not only demonstrates the model’s superior performance—surpassing existing methods by at least 15%—but also highlights the importance of leveraging physical properties.

The significance of this study’s findings extends deep into the field of research. High-accuracy protein classification has implications in biomedical research and pharmaceutical development. By accurately categorizing proteins, researchers can better predict protein functions and interactions, which are crucial for drug discovery and the development of personalized medicine. This capability paves the way for more targeted therapies and faster identification of therapeutic targets, ultimately contributing to advances in treating complex diseases.

Looking to the future, there are several avenues for enhancing this model. First, incorporating more diverse datasets, including those from different organisms and environmental conditions, could improve the model’s generalizability. Secondly, refinement of the model’s architecture to optimize computational efficiency would allow for scaling up to handle larger datasets. Additionally, exploring the integration of newer machine learning techniques, such as reinforcement learning, could provide deeper insights into dynamic protein behaviors and their implications on biological functions.

The continued development and refinement of this model not only promise to elevate the standards of protein classification but also aims to contribute substantially to the broader field of proteomics, where understanding the landscape of proteins and their functions has great potential.

## VI. Availability of Data and Materials

Main database is available at RCSB’s website [23] All code and data available on Github [27].

## VII. Contributions

N.L. and A.K. conceived the review article, collected data, organized figures, and performed all meta-analyses of the literature provided in the paper. C.C. and C.K. contributed to the oversight of this article as co-principal investigators.

## VIII. Acknowledgments

The authors are grateful to the various research studies, clinical trials, and individuals behind the work of documenting protein structures. The authors would like to extend their gratitude to the entire Research and Development team at Revilico for their guidance, support, and contributions to the overall team dynamics.

